# Comparative metabolic profile of mice fed a grain-based controlled diet and a purified controlled diet

**DOI:** 10.64898/2025.12.16.694657

**Authors:** Jenifer Souza de Almeida, Sandra Andreotti, Felipe Nunes de Camargo, Ayumi Cristina Medeiros Komino, Carla Roberta de Oliveira Carvalho

**Affiliations:** Department of Physiology and Biophysics, Institute of Biomedical Sciences, University of São Paulo, São Paulo, Brazil; Department of Physiology, Faculty of Medicine of Ribeirão Preto, University of São Paulo

**Keywords:** Grain-based diet, Purified diet, Metabolic parameters, C57BL/6 male mice

## Abstract

**Purpose:** Grain-based control diets and purified diets are widely used in metabolic studies involving rodents, although their compositional differences may influence physiological outcomes and experimental interpretations. This study aimed to compare the metabolic profile of C57BL/6 mice fed a grain-based diet or a purified diet, both administered under normocaloric conditions.

**Methods:** Male C57BL/6 mice (7 weeks old) were fed a grain-based diet or a purified diet for 12 weeks. Body mass and composition, fasting blood glucose, oral glucose tolerance, and plasma lipid profile (triglycerides, total cholesterol, high-density lipoprotein cholesterol, low-density lipoprotein cholesterol, and very low-density lipoprotein cholesterol) were evaluated. Statistical analyses were conducted using an unpaired Student’s t-test (P ≤ 0.05).

**Results:** No significant differences were observed between groups in the evolution of body mass, body composition, fasting blood glucose, oral glucose tolerance, or plasma lipid parameters.

**Conclusion:** Under normocaloric conditions, grain-based or purified control diets produce equivalent metabolic profiles in C57BL/6 mice, supporting their interchangeable use as control diets in metabolic research studies.

## Introduction

Experimental models of diet-induced obesity are fundamental tools for investigating the metabolic determinants of the disease. However, the choice of control diet used in these models represents a methodological variable that is frequently overlooked. Conventional grain-based diets have a less defined nutritional composition, with intrinsic variations related to processing, plant origin, agricultural conditions, and the presence of bioactive compounds. In contrast, purified diets, such as AIN-93M, have standardized, reproducible formulations that minimize variability between batches [3-6].

The literature indicates that fermentable fibers, non-digestible polysaccharides, and phytochemicals in grain-based diets can modulate the gut microbiota, glycemic homeostasis, and nutrient absorption, thereby influencing relevant metabolic outcomes. Reduced intake of structural dietary components has been associated with a higher risk of insulin resistance, type 2 diabetes mellitus, and metabolic dysfunction-associated steatotic liver disease, reinforcing the importance of the nutritional profile of the diet [11-14].

However, it remains uncertain whether, in the absence of metabolic stress (e.g., high-calorie or high-fat diets), differences between grain-based and purified diets result in measurable metabolic changes. Thus, this study sought to systematically compare the impact of these two formulations on general metabolic parameters in C57BL/6 mice maintained under normocaloric conditions.

## Material and methods

### Animals and experimental design

Seven-week-old male C57BL/6 mice were obtained from the animal facility at the Faculty of Medicine of the University of São Paulo (FMUSP). The animals were kept in specific individual cages, in an environment with controlled temperature (25 ± 2 °C) and a 12-hour light-dark cycle (06:00–18:00), with free access to food and water. After a one-week acclimation period, the mice were randomly allocated into two experimental groups using the Random Sequence generator (https://www.random.org/sequences/): (1) grain-based control diet (CN) (CR-1, Nuvilab, Colombo, PR, Brazil), containing 19% protein, 56% carbohydrates, 3.5% lipids, 5% cellulose and 4.5% vitamins and minerals, totaling 3.20 kcal/g; and (2) purified control diet (CA), formulated according to the recommendation of the American Institute of Nutrition (AIN-93M), composed of 14.1% protein, 75.9% carbohydrates, 10.0% lipids, 5% cellulose and 3.5% vitamins and minerals, with an energy value of 3.6 kcal/g (PragSoluções Biociências, Jaú, SP, Brazil). The animals were fed their respective diets continuously for 12 weeks. The experimental protocol was approved by the Ethics Committee on the Use of Animals of the Institute of Biomedical Sciences of the University of São Paulo (CEUA/ICB-USP), under number 3819110222, on April 26, 2022.

### Oral Glucose Tolerance Test

The oral glucose tolerance test (oGTT) was performed after a 6-hour fasting period. Briefly, a small blood sample was collected from the tail end of the animals by incision to determine baseline blood glucose. Measurements were performed using reagent strips (Roche, Mannheim, Germany) and an Accu-Chek Active glucometer (Roche). Next, the mice received an oral dose (gavage) of 25% glucose solution at 75 mg/100 g body weight. Additional blood samples were collected at 5, 10, 15, 30, 45, 60, and 90 minutes after glucose administration for analysis of plasma glucose concentrations. All determinations were conducted using the same reagent strip system and the Accu-Chek Active glucometer [15–17].

### Body composition

At the end of the experimental period, the body composition of the mice was determined by low-resolution magnetic resonance imaging, using the Minispec LF50 equipment (Bruker, MA, USA), which allows non-invasive quantification of fat and lean mass fractions.

### Euthanasia

After treatment and a 6-hour fasting period, the animals were anesthetized with isoflurane, and euthanasia was performed by rapid decapitation. The collected blood was immediately placed on ice and stored at −80 °C until the biochemical analyses were performed [18].

### Biochemical Analysis

Initially, blood was collected and centrifuged (4 °C, 4,000 × g, 30 min). Then plasma concentrations of triglycerides (TG), total cholesterol (TC), and high-density lipoprotein (HDL-C) were determined using commercial enzyme kits, according to the manufacturer’s instructions (TG: Cat. No. 87; TC: Cat. No. 76; HDL-C: Cat. No. 145; LabTest, Belo Horizonte, MG, Brazil). Very low-density lipoprotein (VLDL-C) and low-density lipoprotein (LDL-C) fractions were calculated according to a previously described methodology [19].

### Data Analysis

Data normality was verified using the Shapiro–Wilk test, while homoscedasticity was assessed using the Brown–Forsythe test. Depending on the data distribution, analyses were conducted using the unpaired Student’s t-test or, when appropriate, the corresponding nonparametric test. The significance level adopted was 5% (P ≤ 0.05).

## Results

### Body composition and body mass are not affected by the type of control diet

Animals fed both types of diet started the dietary protocol with similar body masses (Figure 1, a - CN: 21.480 ± 0.576 vs. CA: 22.760 ± 1.322 g; P = 0.0825), indicating adequate initial randomization. Throughout the 12-week intervention, weekly body mass monitoring showed a pattern of progressive and similar body mass gain between groups. Analysis of the area under the curve (AUC) of body mass (Figure 1, b, and c - CN: 265.060 ± 10.494 vs. CA: 287.620 ± 26.843 g.week; P = 0.1182) confirmed that the type of control diet did not exert a significant effect on body mass throughout the experimental period. Additionally, the assessment of final body mass (Figure 1, d - CN: 24.600 ± 2.191 vs. CA: 26.848 ± 3.296 g; P = 0.2398) revealed no statistically significant difference between the groups. Furthermore, body composition, assessed by magnetic resonance imaging, showed that both lean mass content (Figure 1, e - CN: 17.148 ± 1.294 vs. CA: 19.369 ± 3.008 g; P = 0.4206) and fat mass content (Figure 1, f - CN: 1.566 ± 0.754 vs. CA: 1.954 ± 0.509 g; P = 0.3671) remained unchanged between the experimental groups.

**Fig. 1.**
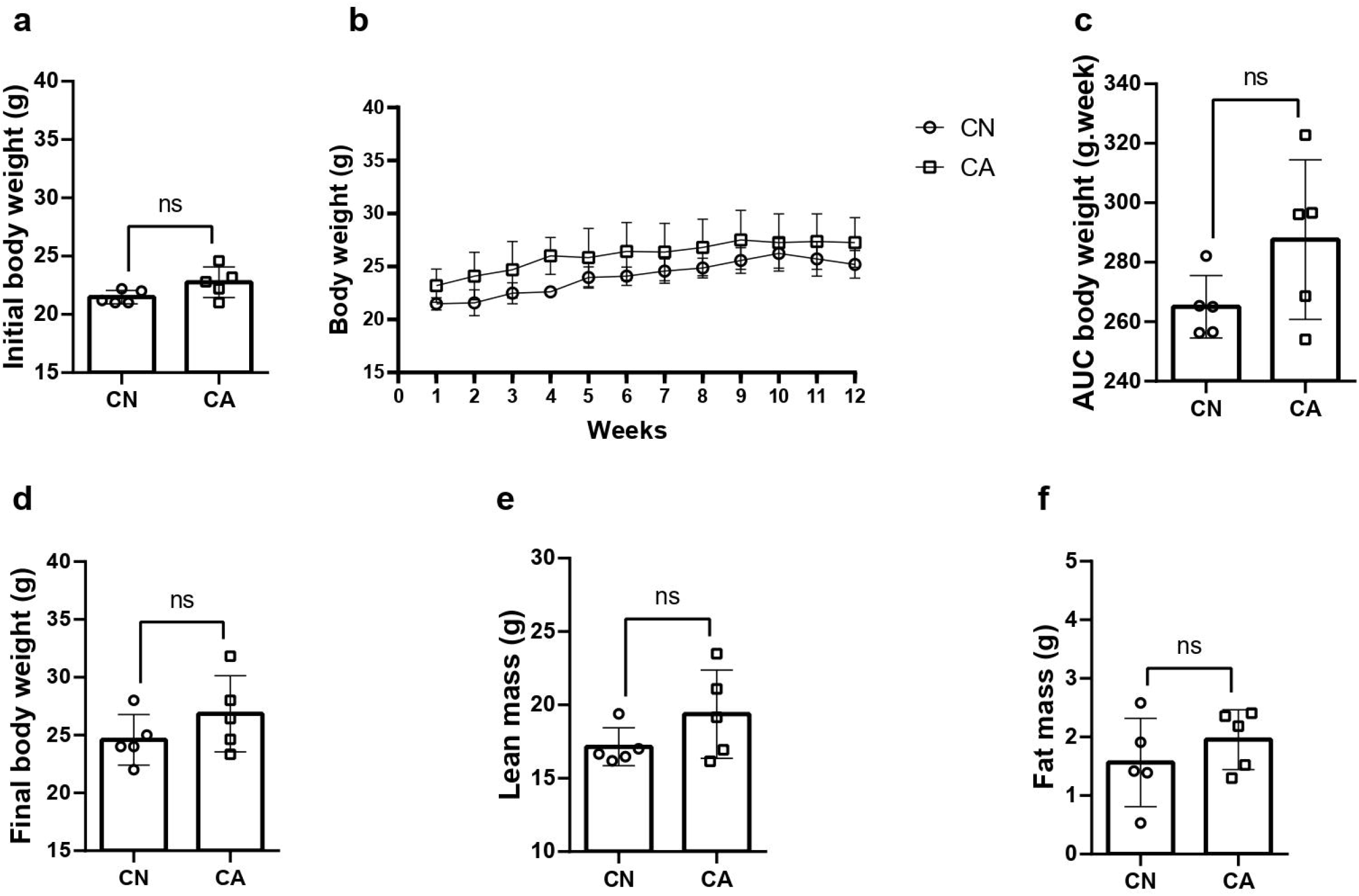
Body mass and composition. (a) Initial body mass (n = 5 in both groups); (b) Time course throughout the dietary protocol (n = 5 in both groups); (c) AUC of body mass throughout the dietary period (n = 5 in both groups); (d) Final body mass (n = 5 in both groups); (e) Lean mass (n = 5 in both groups); (f) Fat mass (n = 5 in both groups). Results were obtained using Student’s t-test and are presented as mean ± standard deviation (SD), with a significance level of P ≤ 0.05. Each scatter plot indicates a sample unit, and “ns” indicates the absence of a statistically significant difference between groups.

These findings indicate that the choice between a grain-based and a purified control diet did not significantly affect weight dynamics or body composition in C57BL/6 mice throughout the experimental period. Thus, under normocaloric conditions, both formulations are equally capable of supporting growth and maintaining body mass without promoting significant differences in adipose tissue accumulation or lean mass.

### The type of control diet does not affect fasting blood glucose or glucose tolerance

The data in Figure 2 indicate that, consistent with the findings on body mass and body composition, the type of control diet did not significantly affect the animals’ glycemic parameters. Fasting blood glucose (Figure 2, a - CN: 7.012 ± 0.792 vs. CA: 7.844 ± 1.464 mmol·L□^1^; P = 0.2959) remained statistically similar between the groups fed a grain-based diet and a purified diet.

**Fig. 2.**
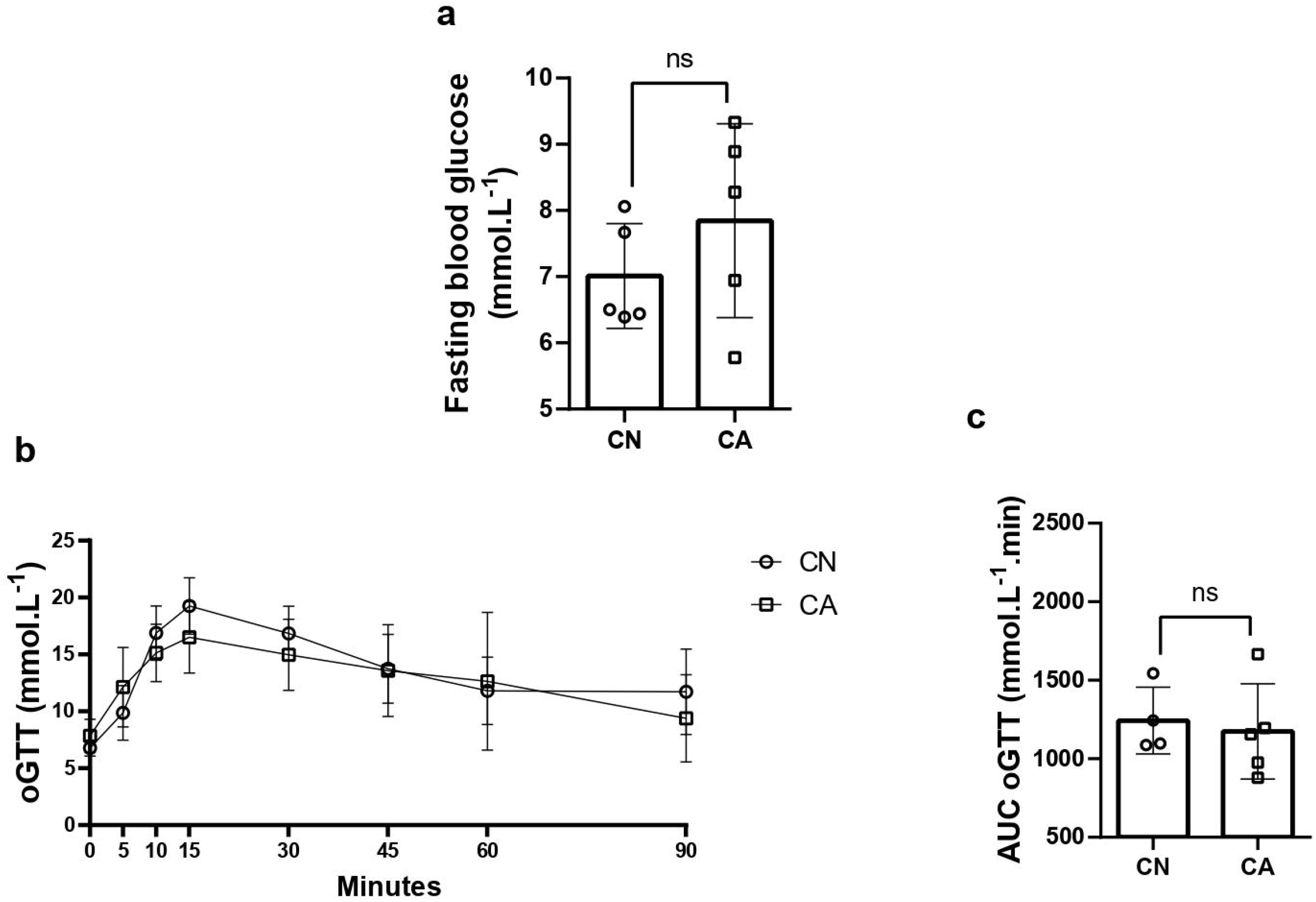
Fasting blood glucose and oral glucose tolerance test. (a) Fasting blood glucose (n = 5 in all groups); (b) Time course of the oral glucose tolerance test (n = 4 in the CN group, and n = 5 in the CA group); (c) AUC of the oral glucose tolerance test (n = 4 in the CN group, and n = 5 in the CA group). Results were obtained using Student’s t-test and are presented as mean ± standard deviation (SD), with a significance level of P ≤ 0.05. Each scatter plot indicates a sample unit, and “ns” indicates the absence of a statistically significant difference between groups.

Additionally, the OGTT assessment revealed no significant differences between the experimental groups. Analysis of the oGTT AUC (Figure 2, b and c - CN: 1243.250 ± 212.887 vs. CA: 1174.440 ± 303.679 mmol·L□^1^·min; P = 0.7138) confirmed that the ability to metabolize glucose was equivalent in both groups, suggesting that the choice between a grain-based control diet or a purified diet does not significantly impact basal glycemic homeostasis or the response to glucose overload in C57BL/6 mice.

### The type of control diet does not alter the plasma lipid profile

Analysis of the animals’ plasma lipids after the experimental period revealed that the type of control diet did not generate significant differences in the parameters evaluated. For example, TG, TC, HDL-C, LDL-C, and VLDL-C levels did not differ significantly between animals fed grain-based feed and those fed a purified diet (Table 1).

**Table 1.**
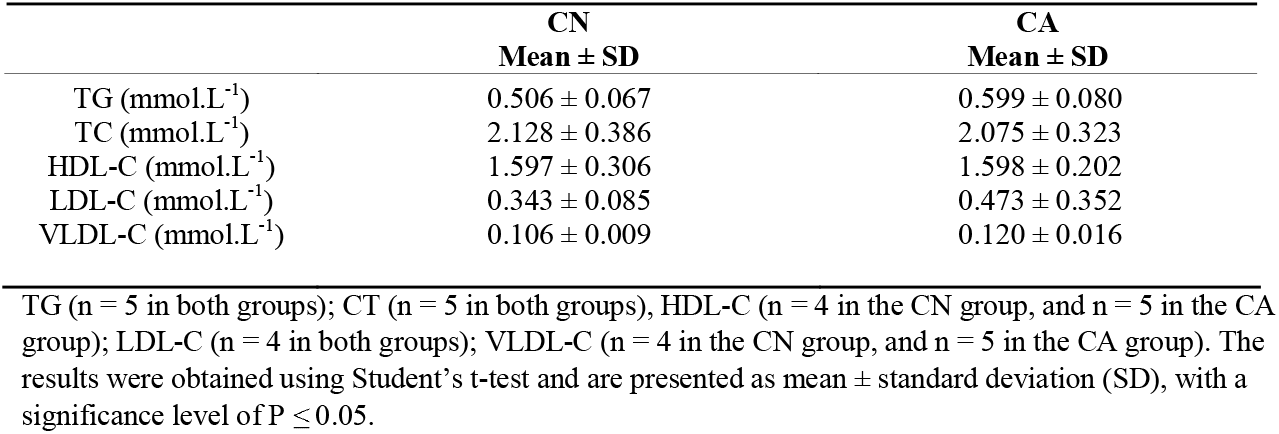
Lipid Profile.

| CN | CA |
| --- | --- |
| Mean ± SD | Mean ± SD |

|  |  |  |
| --- | --- | --- |
| TG (mmol.L<sup>-1</sup>) | 0.506 ± 0.067 | 0.599 ± 0.080 |
| TC (mmol.L<sup>-1</sup>) | 2.128 ± 0.386 | 2.075 ± 0.323 |
| HDL-C (mmol.L<sup>-1</sup>) | 1.597 ± 0.306 | 1.598 ± 0.202 |
| LDL-C (mmol.L<sup>-1</sup>) | 0.343 ± 0.085 | 0.473 ± 0.352 |
| VLDL-C (mmol.L<sup>-1</sup>) | 0.106 ± 0.009 | 0.120 ± 0.016 |

TG (n = 5 in both groups); CT (n = 5 in both groups), HDL-C (n = 4 in the CN group, and n = 5 in the CA group); LDL-C (n = 4 in both groups); VLDL-C (n = 4 in the CN group, and n = 5 in the CA group). The results were obtained using Student's t-test and are presented as mean ± standard deviation (SD), with a significance level of $P \leq 0.05$ .

These findings indicate that during the evaluated intervention period, the choice between a grain-based control diet and a purified diet does not significantly affect plasma lipids, suggesting that both formulations are equivalent in maintaining the lipid profile in the experimental model used.

## Discussion

This study evaluated the impact of a grain-based control diet and a purified AIN-93M diet on general metabolic parameters in C57BL/6 mice maintained under normocaloric conditions. The results revealed no significant differences between the groups in terms of body mass, body composition, fasting blood glucose, glucose tolerance, or plasma lipid profile.

The findings regarding body mass and body composition indicate that, in the absence of caloric overload, both dietary formulations are equally capable of supporting growth and maintaining lean and fat mass without inducing detectable phenotypic changes. This absence of effects is consistent with previous studies demonstrating that, when there are no obesogenic stimuli or excess lipids, the composition of the control diet exerts a limited influence on weight gain and the distribution of body compartments [5,20,21].

Similarly, glycemic parameters remained stable between groups, suggesting that moderate differences in nutritional composition, such as variations in starch content, fiber, or energy density, were insufficient to alter glucose homeostasis over the 12-week intervention. The stability observed in the oGTT reinforces the notion that carbohydrate metabolism in C57BL/6 mice exhibits a high capacity for adaptation, being more sensitive to intense nutritional challenges, such as high-fat or high-calorie diets, than to subtle adjustments in the composition of control diets [20,22].

The similarity in plasma lipid profile also demonstrates this metabolic robustness. Concentrations of TG, TC, HDL-C, LDL-C, and VLDL-C remained statistically equivalent, indicating that, under balanced caloric intake, the choice between a grain-based and a purified diet does not significantly affect lipid metabolism. Although grain diets have higher fiber and bioactive compound content, these characteristics did not translate into detectable phenotypic changes. It is plausible that a challenging metabolic environment, such as physiological stress, inflammation, or excessive lipid supply, which could have potentiated differences between the formulations, was absent [20,21].

Taken together, the results suggest that homeostatic and metabolic compensation mechanisms played a central role in maintaining a similar phenotype between the groups. C57BL/6 mice exhibit high metabolic flexibility, adjusting voluntary intake, feed efficiency, metabolic rate, substrate oxidation, and even fecal energy excretion according to moderate dietary variations. These mechanisms may have allowed the animals to balance the nutritional differences between the two formulations, thereby avoiding detectable systemic alterations over the 12-week intervention.

Several limitations should be considered when interpreting the results. For example, daily food intake, energy expenditure, and metabolic efficiency were not directly assessed; these variables are essential for elucidating the compensatory mechanisms that may help maintain homeostasis. Moreover, specific insulin sensitivity tests, which could detect more subtle changes in glycemic dynamics not revealed by the oGTT, were not performed. Similarly, molecular analyses, including gene or protein expression in metabolically relevant tissues such as liver, adipose tissue, muscle, and kidney, were not conducted, limiting the understanding of the underlying cellular adjustments. Lastly, although the sample size is adequate to detect moderate effects, small-magnitude metabolic differences may have gone unnoticed.

The results presented have relevant implications for studies using control diets in rodents. The absence of metabolic differences between the grain diet and the purified diet indicates that, under normocaloric conditions and without obesogenic challenges, both can serve as equivalent controls in experiments whose objective is not to induce systemic phenotypic alterations but rather to ensure adequate nutrition. This equivalence is particularly important for laboratories seeking experimental standardization, since purified diets exhibit less variability between batches, while grain diets offer lower cost and greater availability. Furthermore, the findings reinforce that detectable metabolic modifications in C57BL/6 mice generally require more intensive nutritional interventions, such as high-fat, high-calorie diets or specific supplements, highlighting the importance of aligning the diet type with the experimental objective.

## Statements and Declarations

## Acknowledgments

We want to thank the National Council for Scientific and Technological Development (CNPq, grant no. 140217/2022-3) and the São Paulo Research Foundation (FAPESP, grants no. 2016/25129-4, 2017/18972-0, and 2021/10469-2) for the financial support. We would also like to thank Gustavo Cândido e Cruz, M.Sc.; Beatriz Garcia de Carvalho, M.Sc.; Sandro Leão Matos, Ph.D.; and Andressa Godoy Amaral, Ph.D.,for their initial support during the experimental phase.

## Author Contributions

Jenifer Souza de Almeida and Carla Roberta de Oliveira Carvalho contributed to the conception and design of the study. Jenifer Souza de Almeida, Sandra Andreotti, Felipe Nunes de Camargo and Ayumi Cristina Medeiros Komino collected the data. Jenifer Souza de Almeida prepared the materials, analyzed the data and wrote the first draft of the manuscript, and all authors commented on previous versions. All authors read and approved the final manuscript.

## Ethics approval

The Ethics Committee on the Use of Animals of the Institute of Biomedical Sciences of USP (CEUA/ICB-USP) approved the use of animals in this study on April 26, 2022, under protocol number 3819110222.

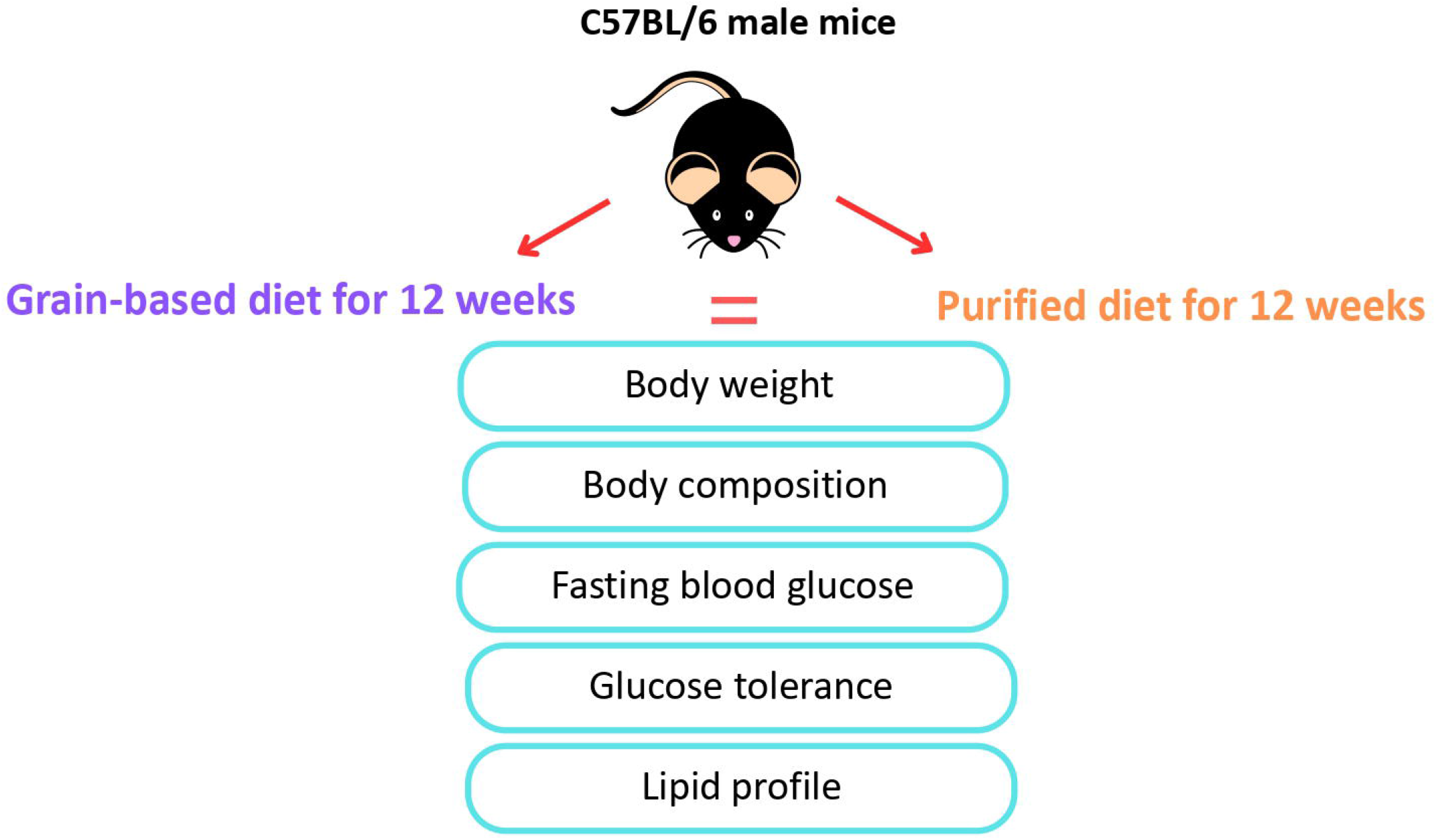

